# A role for Myosin in triggering and executing amnioserosa cell delaminations during dorsal closure

**DOI:** 10.1101/2025.03.21.644291

**Authors:** Nicole Gorfinkiel, Yanara Ferrer, Jon Recalde, Javier Gutiérrez, Guillermo Sáez

## Abstract

The remodeling of epithelial tissues is a critical process in morphogenesis, often involving the apoptotic removal of individual cells while preserving tissue integrity. In *Drosophila*, the amnioserosa—a highly dynamic extra-embryonic tissue—undergoes extensive remodeling, culminating in its complete elimination at the end of dorsal closure. While apoptotic cell delaminations in the amnioserosa have been proposed to contribute to dorsal closure, the cellular mechanisms underlying this process remain poorly understood. In this study, we have investigated actomyosin dynamics during cell delaminations and analyzed the consequences of perturbing non-muscle Myosin activity globally in the entire tissue as well as locally in groups of cells. We found that Myosin plays an essential role in both triggering and executing cell delaminations, with high Myosin contractility promoting cell delamination via caspase activation. Additionally, our results suggest that cell delaminations are governed by both cell-autonomous Myosin dynamics and mechanical cues from the tissue environment. Together, these findings provide new insights into the regulation of epithelial cell removal and the complex interplay between apoptotic and mechanical signals during tissue remodeling.

## INTRODUCTION

The remodelling of epithelial tissues is a fundamental process in morphogenesis, giving rise to a diverse array of structures, including sheets, tubes, and cysts, which are essential for organ formation and function. A key aspect of epithelial remodelling is the apoptotic removal of individual cells from the epithelium while maintaining overall tissue integrity. Apoptotic cell elimination has been observed in diverse contexts contributing both to tissue morphogenesis and to the maintenance of tissue homeostasis^1–3^. In some instances, entire epithelial tissues are removed and replaced, as occurs during embryogenesis in the extra-embryonic tissues of insects^4^ and during metamorphosis^5^.

The amnioserosa is the single extra-embryonic tissue present in Drosophila^6^. It is specified during early embryonic development by the dorso-ventral patterning system and plays essential functions during two vital morphogenetic movements, germband retraction and dorsal closure^7^. Amnioserosa cells undergo dramatic cell shape changes during embryogenesis. During germband elongation, these cells elongate and transition from a columnar to a squamous morphology through a process known as rotary cell elongation^8^. As development progresses into germband retraction, amnioserosa cells shorten, become isodiametric, and exert a pulling force that contributes to germband retraction^9,10^. Finally, during dorsal closure, amnioserosa cells exhibit dynamic fluctuations in apical surface area, driven by periodic contractions of the actomyosin cytoskeleton, and progressively reduce their apical surface^11–14^. Around 10% of the cells of the tissue basally delaminate before closure is completed^15–18^. These cellular processes, along with a decrease in volume^19^, generate a morphogenetic force that drives the movement of the lateral epidermal sheets towards the dorsal midline. Ultimately, the entire amnioserosa tissue is internalized and degenerates by apoptosis^20–22^.

Amnioserosa cell delaminations are apoptotic, and although they are not essential for closure, enhancing or supressing apoptosis increases or reduces cell delaminations respectively, leading to either an acceleration or a slowing down of closure^16^. This led to the suggestion that apoptotic cell delaminations in the amnioserosa provides an apoptotic force that contributes to closure. Additionally, inhibiting apoptosis in the amnioserosa also prevents cell volume decrease in the bulk of the tissue^19^, showing that the apoptotic pathway controls both individual delaminations and the remodelling of the entire tissue.

The cellular mechanisms underlying the elimination of cells from an epithelium have been studied in several tissues, revealing the involvement of both cell autonomous and non-autonomous mechanisms. In the apoptotic cell, either an actomyosin ring^23–26^ or medioapical actomyosin contractions^27–29^ can contribute to autonomous-cell constriction, while neighbouring cells form a supracellular actomyosin cable that aids in the expulsion of the cell from the tissue^23,24,26^. In the amnioserosa, high levels of reactive oxygen species induce caspase activity and cell delaminations through the reorganization of non-muscle Myosin distribution^18^. During the early stages of dorsal closure, delaminating cells transition abruptly from a pulsatile behaviour to a rapidly, unpulsed, contracting behaviour^30^. This delamination transiently affects the oscillating behaviour of the neighbouring cells, and changes in Myosin dynamics have been observed in both the delaminating cell and its neighbours^30^. However, the precise cellular mechanisms underlying cell delamination in the amnioserosa remain unclear. Additionally, mechanical stresses have been proposed as potential triggers for both individual cell delamination of cells and the degeneration of the entire amnioserosa, but evidence for this has been lacking. In this work, we have investigated the cellular mechanisms driving cell delaminations in the amnioserosa and analyzed the consequences of perturbing non-muscle Myosin activity globally in the entire tissue as well as locally in groups of cells. Our results indicate that Myosin plays an active role in both the initiation and execution of cell delaminations, and show that cells with high contractility can undergo delamination through the induction of caspase activity.

## RESULTS

### 1) Actomyosin dynamics during amnioserosa cell delaminations

Cell delaminations in the amnioserosa can be observed from the end of germband retraction until the completion of dorsal closure. Although their number varies among embryos, they exhibit a consistent spatial pattern, occurring mostly in the anterior half of the amnioserosa and at the posterior canthi^15,17,18^ (see below).

In epithelial tissues, cell delaminations are usually mediated by T2 transitions, in which multiple vertices converge to form a single vertex, generating a rosette-like structure^23,31^. In the amnioserosa (Movie 1), while some cells delaminate through T2 transitions (Figure 1A, Á, A”), we also observe delaminating cells that do not form rosettes. Instead, these cells leave a new junction once they are removed from the tissue. This type of delamination occurs in anisotropic cells where the longer junctions, usually dorso-ventrally oriented, approach and fuse, forming a new junction between new neighbours (Figure 1B, B’, B”, Movie 1B). The frequency of both types of delamination is variable among embryos, with a slightly higher average number of delaminations forming rosettes than forming a new junction, but these differences are not statistically significant (Figure S1). We found that cells leaving a rosette are more isotropic at the onset of delamination than cells leaving a junction (Figure S1), suggesting that cell shape determines the geometry of the delamination process. The ectopic expression of Diap1 (Drosophila inhibitor of apoptosis 1), p35 (baculovirus apoptosis inhibitor) or RHG miRNA (three microRNAs that simultaneously inhibit the proapoptotic genes *reaper, hid* and *grim*^32^) in the amnioserosa prevent all cell delaminations (see^15,16,18^ and Movie 2).

**Figure 1.**
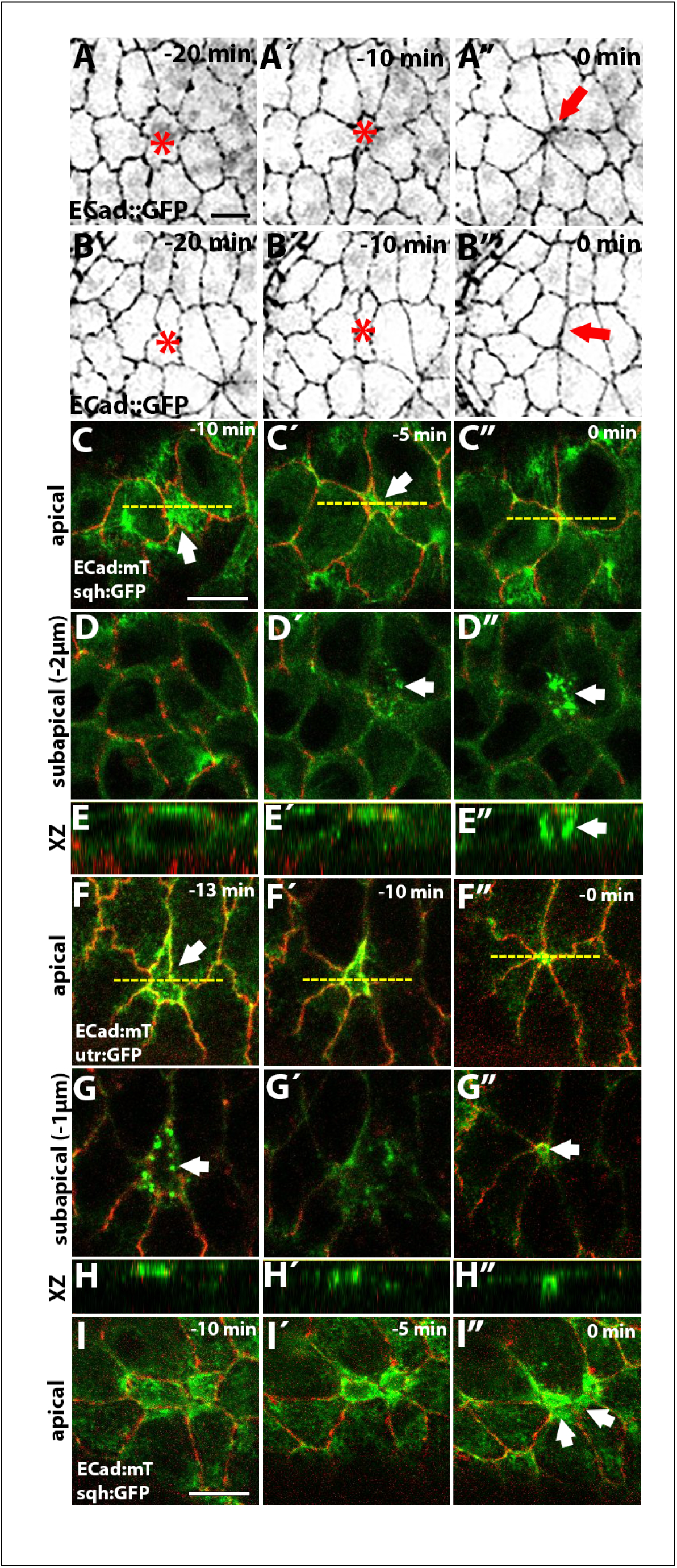
Actomyosin dynamics during amnioserosa cell delamination. (A,B) Still images from a time-lapse movie (Movie 1) of a dorsal closure ECad:GFP embryo at different time points before the completion of cell delamination (asterisk), showing the formation of a rosette-like structure (A) or the formation of a new junction (B, arrow, see also Movie 1B). (C-E) Still images from a time-lapse movie of an early dorsal closure sqh:GFP; ECad:mT embryo (C-E, Movie 3) or ECad:mT/ utr:GFP (F-H, Movie 4) showing an apical (C, F) and subapical (D, G) planes, as well as an orthogonal view (E, H) at indicated times before the completion of cell delamination. Panels C, D and G are individual z-sections, while panels F, due to the curvature of the tissue where delamination occurs, are the maximum intensity projection of the 2 planes with the brightest ECad signal. Notice the stabilisation of medioapical actomyosin during the apical-contraction phase (C, Ć, E, É, F, F’, H, H’), the appearance of actin (G, arrow) and MyoII (D’, D”, arrow) puncta, and the formation of an actin ring during the final phase of delamination (G”, arrow). The yellow dotted lines in C and F indicate the position of the orthogonal view shown in E and H. These are representative examples of early delaminating cells, based on the analysis of 12 cells from 7 embryos (sqhGFP; ECad:mT) and 10 cells from 3 embryos (ECad:mT/utr:GFP). (I-I”) Still images from a time-lapse movie of a late dorsal closure sqh:GFP; ECad:mT embryo. Notice the accumulation of Myosin in neighbouring cells (I”, arrows), which coincides with a decrease of ECad:mT at cell junctions. This is a representative example of a late delaminating cell based on the analysis of 14 cells from 3 embryos. Scale bar represents 10µm in this and all subsequent figures (if a panel lacks a scale bar, it is the same as in the preceding panel).

It has been reported that cell delaminations in different tissues can be driven either by a cortical actomyosin ring or by pulses of contractile medioapical actomyosin^24,26–29^. To investigate the mechanisms driving cell delaminations in the amnioserosa, we visualized non-muscle Myosin (MyoII) and actin distribution in delaminating cells from live embryos carrying the ECad:mTomato (ECad:mT) and the sqh:GFP knock-in alleles (Figure 1C-E”, I-I”; Movie 3), or ECad:mT and the actin reporter utrophin fused to GFP (utr:GFP) (Figure 1F-H”; Movie 4). Since MyoII subcellular localization and dynamics changes as dorsal closure progresses, characterized by increasing junctional MyoII and stabilisation of the medioapical pool^12,13^, we examined cell delaminations at both early and late stages. During early stages, as the delaminating cell begins to apically constrict, we observed a stabilisation of medioapical MyoII (Figure 1C, Ć, E, É) as well as actin (Figure 1F, F’, H, H’), with no apparent accumulation of junctional Myosin nor actin. Early delaminating cells are pulsatile^11–14^, and the onset of apical constriction coincides with the stabilisation of a medial MyoII pulse, giving rise to a non-pulsatile delaminating cell, as previously observed^30^. As the apical cell area reduces, cell junctions show an inward curvature (Figure 1C, Ć, arrow), suggesting the existence of a pulling force from the medial part of the delaminating cell. Interestingly, we observed the appearance of actin puncta that start to form subapically (1-2 µm basal to the plane of ECad brightest signal) (Figure 1G and S2). MyoII puncta are also observed although they appear later than the actin puncta (Figure 1D’). As the apical cell area reduces, actin puncta can be seen to re-localize apically at the level of ECad (Figure 1G’ and Figure S2 for a more detailed description of actin dynamics during the delamination process) and eventually shows a continuous junctional accumulation (Figure 1F’). Once the cell ingresses, an actin junctional ring is observed 1 µm underneath the plane of the epithelium (Figure 1F”, G”, H”), while MyoII still retains a punctate pattern (Figure 1C”, D”, E”). Such cytoskeletal dynamics has been observed in both isotropic and anisotropic cell delaminations (Figure S3).

Since MyoII subcellular localization and dynamics changes as dorsal closure progresses, characterized by increasing junctional MyoII and stabilisation of the medioapical pool^12,13^, we also examined cell delaminations during late dorsal closure stage (Figure 1I, Í, I”). Late delaminating cells also show a stabilisation of medioapical MyoII that spans the whole apical cortex as the cell area reduces (Figure 1I). An actomyosin ring is also observed as the delamination process proceeds (Figure 1I’, I”). In contrast to what has been observed in other epithelia, we did not observe the formation of a supracellular actomyosin ring in neighbouring cells. However, in some late delaminating cells, MyoII accumulated in neighbouring cells at the junctions with the delaminating cell (Figure 1I”, arrows) coincident with a decrease in ECad levels (Figure S4). We propose that this could be a mechanism to maintain junction integrity during the delamination process^33^ but would not be required to expulse the cell from the tissue.

Altogether, these observations indicate that amnioserosa cells delamination involves the stabilisation of medioapical Myosin and the formation of an actomyosin ring as delamination progresses, with no evidence for an active contribution of neighbouring cells to the process.

The Rho family of GTPases is a central regulator of F-actin, MyoII activity and contractility^34^. Both Rho-dependent and independent mechanisms for apoptotic cell elimination have been reported^35^. In the amnioserosa, using a Rho live sensor consisting of the Rho1 GTP-binding domain of anillin to GFP (aniRBD:GFP)^36^, we have observed an increase in medioapical Rho activity as soon as the apical cell area of a delaminating cell begins to decrease (Figure 2B, B’, B”, Movie 5). Thus, it is possible that apoptotic signals induce medioapical Rho activity, which in turn regulates the cytoskeletal changes described above contributing to the execution of the delamination process. Interestingly, during late stages of dorsal closure, all amnioserosa cells showed medioapical accumulation of the Rho sensor (Figure 2A, Á). Since medioapical Rho activity is observed in apoptotic delaminating cells, and caspase activity has been detected throughout the amnioserosa tissue during late dorsal closure stages^19,37,38^ (and see below), we asked whether this late medioapical Rho accumulation depends on caspase activity. The ectopic expression of Diap1 in the amnioserosa, while preventing cell delamination, did not prevent the medioapical accumulation of the Rho sensor in the bulk of the tissue during late dorsal closure (Figure 2C). This result suggests that Rho activity at the medioapical cortex of amnioserosa cells is regulated differently in delaminating cells versus the bulk of the tissue, where it appears to be independent of apoptotic signals.

**Figure 2.**
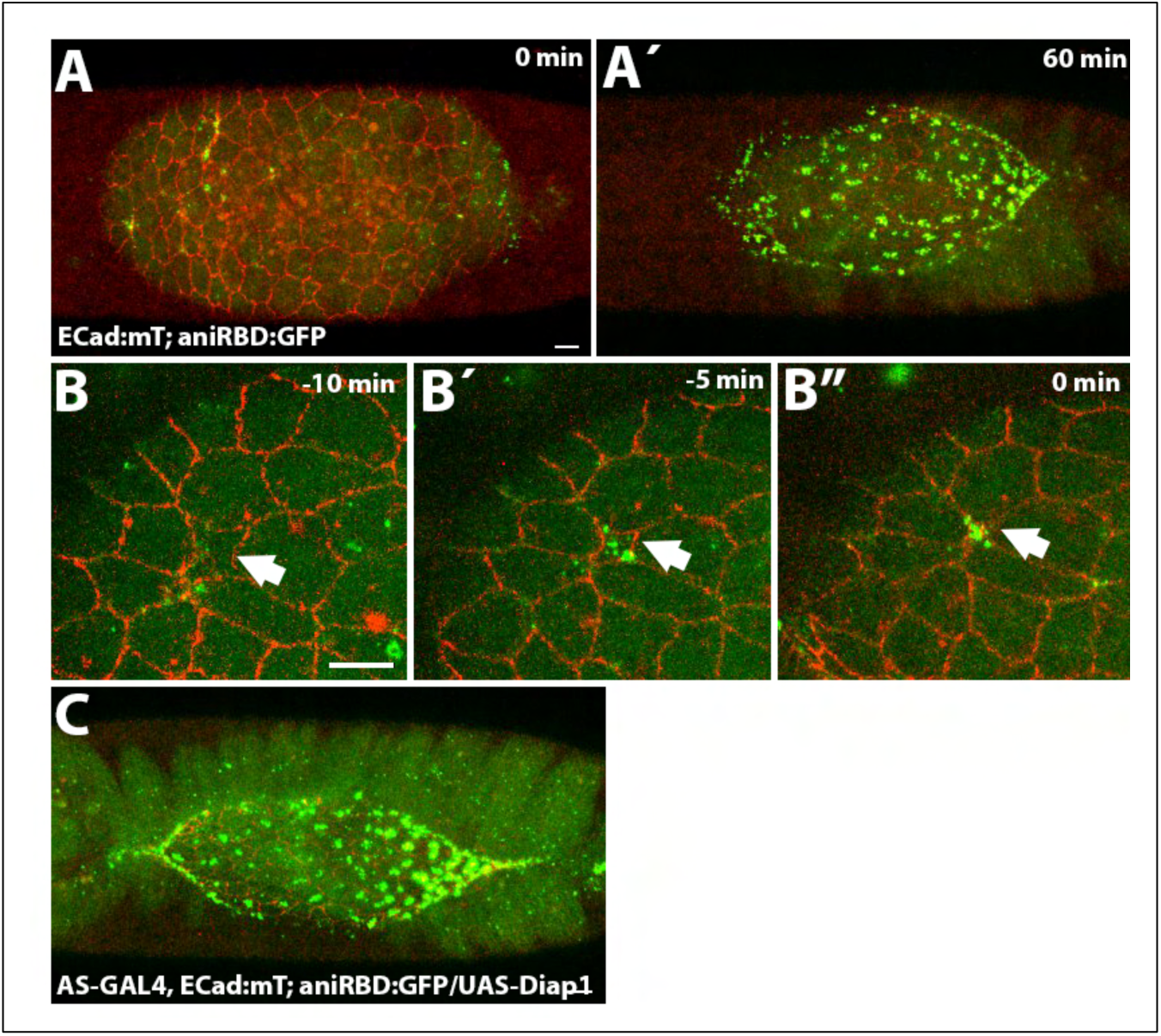
Rho activity in the amnioserosa. (A, Á) Still images from a time-lapse movie of a dorsal closure Drosophila embryo carrying ECad:mT and the Rho sensor (aniRBD:GFP). During most of dorsal closure, the activity of the Rho sensor is only detected in delaminating cells (A). By the end of the process, most of the cells show medioapical accumulation of the Rho sensor (Á), in 9 out of 14 embryos analyzed. (B) Magnification of the same time-lapse than in A (Movie 5), showing a delaminating cell (arrows) and the medioapical accumulation of the Rho sensor at indicated time points before the completion of delamination. This is a representative example of a delaminating cell based on the analysis of 10 cells from 6 embryos. (C) Still image from a time-lapse movie of a late dorsal closure embryo ectopically expressing Diap1 in the amnioserosa showing medioapical accumulation of the Rho sensor (n=3).

### 2) Perturbing Myosin levels in the whole tissue does not alter the number of cell delaminations

To investigate the requirement for Myosin in the initiation of cell delaminations and in the elimination of cells from the tissue, we aimed to perturb Myosin II activity in the whole tissue. We have previously shown that the ectopic expression in the amnioserosa of a constitutive active form of Mbs, MbsN300^39^ (Mbs is the Myosin binding subunit of the protein phosphatase PP1), decreases phosphorylated Myosin, decreases the rate of contraction of amnioserosa cells and slows dorsal closure^40,41^. Here, we analyzed whether cell delaminations are affected upon ectopic expression of MbsN300 (Figure 3A-D).

**Figure 3.**
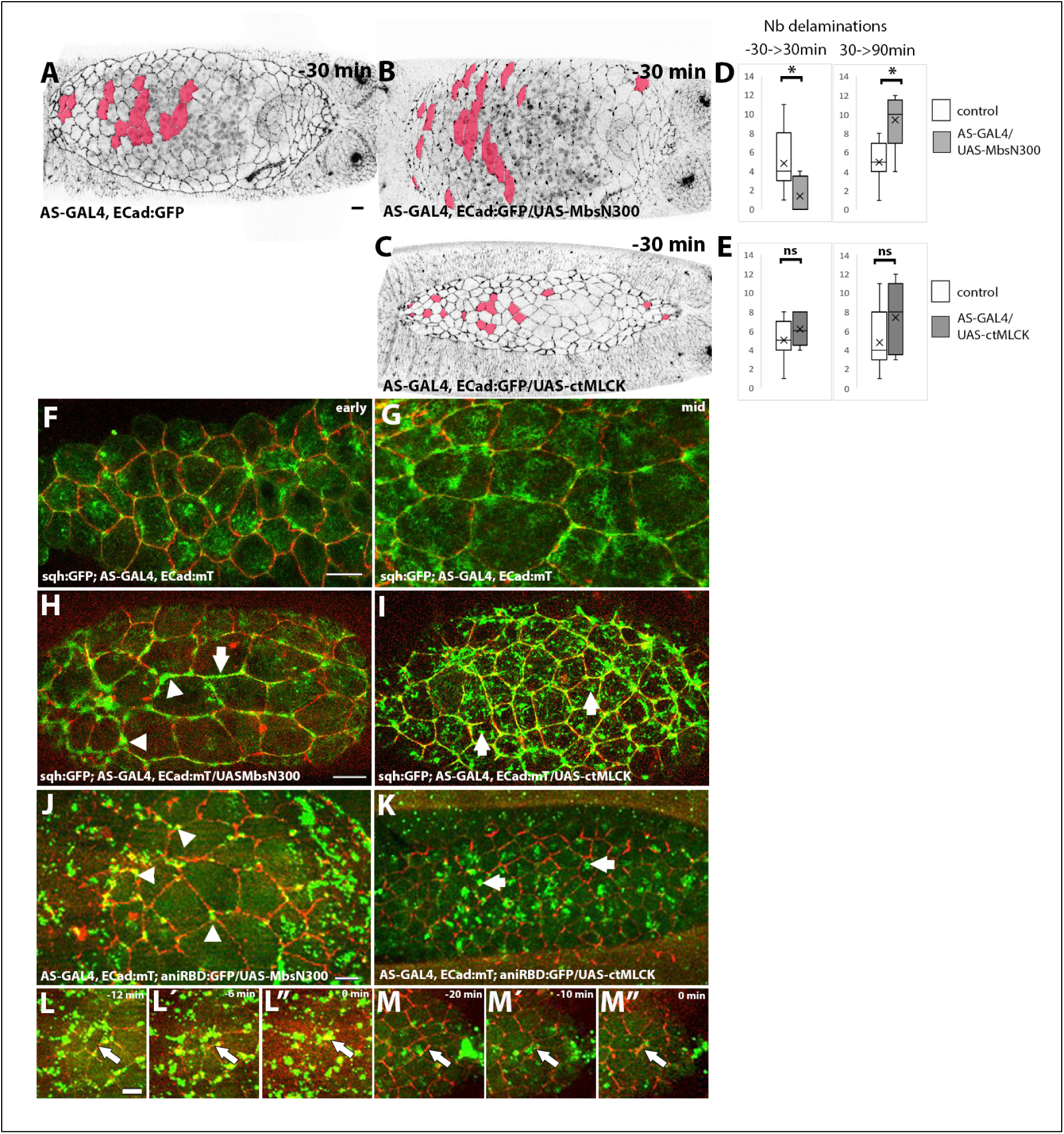
Effect of perturbing Myosin activity in the amnioserosa on cell delaminations and actomyosin contractility. (A-C) Still image of a control ECad:GFP embryo (A, Movie 1), an AS-GAL4, ECad:GFP/UAS-MbsN300 (B, Movie 6) and an AS-GAL4, ECad:GFP/UAS-ctMLCK embryo (C, Movie 7) at -30 min of dorsal closure (staging relative to the distance of posterior spiracles). The cells labelled in red are cells that delaminate during dorsal closure. (D) Number of delaminating cells in AS-GAL4, ECadGFP embryos (n=11) and AS-GAL4, ECadGFP/UAS-MbsN300 embryos (n=5), between -30 and 30 min (left panel, p = 0.039, Wilcoxon-Mann-Whitney test) and between 30 and 90 min (right panel, p = 0.029, Wilcoxon-Mann-Whitney test). Although the total number of delaminating cells is similar between the two genotypes, AS-GAL4, ECadGFP/UAS-MbsN300 embryos show a delay in the appearance of cell delaminations, with most of them occurring from 30 min of dorsal closure. (E) Number of delaminating cells in AS-GAL4, ECadGFP (n=11) and AS-GAL4, ECadGFP/UAS-ctMLCK (n=5) embryos, between -30 and 30 min (left panel) and between 30 and 90 min (right panel). No significant differences in the total number of delaminating cells nor a change in the timing of appearance of cell delaminations is observed in these embryos (Wilcoxon-Mann-Whitney test) (F, G) Still images from time-lapse movies of example sqh:GFP; ECad:mT embryos at early (F) and mid dorsal closure (G). (H, I) Still images from time-lapse movies of example sqh:GFP; AS-GAL4, ECad:mT/UAS-MbsN300 (H) and sqh:GFP; AS-GAL4, ECad:mT/UAS-ctMLCK (I) embryos (n=4 and n=3, respectively). (J, K) Still images from time-lapse movies of AS-GAL4, ECad:mT; aniRBD:GFP/UAS-MbsN300 (J, Movie 8) and AS-GAL4, ECad:mT; aniRBD:GFP/UAS-ctMLCK (K, Movie 9) embryos (n=4 and n=5, respectively). Notice the accumulation of Sqh:GFP and the Rho sensor at bicellular and tricellular junctions in H, J (arrow and arrowheads respectively) and their medioapical accumulation in I, K (arrows). (L, M) Still images of example delaminating cells from AS-GAL4, ECad:mT; aniRBD:GFP/UAS-MbsN300 (L) and AS-GAL4, ECad:mT; aniRBD:GFP/UAS-ctMLCK (M) embryos at indicated time points before the completion of delamination. These are example cells based on the analysis of 5 (MbsN300) and 8 (ctMLCK) delaminating cells from 3 different embryos for each genotype.

We are interested in analyzing the contribution of Myosin activity in triggering cell delaminations. To this end, we compared the number of delaminations between the two genotypes occurring during the same time window, in order to avoid variations due to differences in the duration of dorsal closure. Specifically, we only considered cell delamination occurring between -30 min to 100 min of dorsal closure, staging the embryos according to the position of the posterior spiracles (see Materials and Methods). Surprisingly, we found that the total number of cell delaminations in MbsN300 embryos in this time window is indistinguishable from that in control embryos. However, when comparing the timing of cell delaminations, we observed a delay: the number of cell delaminations between -30 and 30 min of dorsal closure is lower in AS-GAL4, ECad:GFP/UAS-MbsN300 than in controls but is higher between 30 and 100 min of dorsal closure (Figure 3A, B, D, Movie 6). These results suggest an early requirement of MyoII for triggering cell delaminations, however, unexpectedly, during later stages, cell delaminations resume and become more frequent.

To explore what could be driving late delaminations in these embryos, we analyzed MyoII localization in sqh:GFP; AS-GAL4, ECad:mT/UAS-MbsN300 embryos. While MyoII localizes in medioapical foci in early dorsal closure, in older embryos, MyoII shows an aberrant distribution, accumulating at bicellular junctions and cell vertices, without forming the dense medioapical mesh observed in control embryos (Figure 3F, G, H, see also^33^). Interestingly, the Rho sensor is also mislocalized, accumulating at cell junctions and cell vertices (Figure 3J). In these embryos, delaminating cells display junctional rather than medioapical Rho1 activity (Figure 3L, Ĺ, L”, Movie 8), along with strong junctional MyoII (Figure S5).

Reciprocally, we aimed to increase Myosin activity in the amnioserosa by ectopically expressing a constitutive active form of Myosin Light Chain Kinase (ctMLCK). We have previously shown that these cells are more contracted and accumulate both junctional and medioapical MyoII^12,42^. While in other tissues increasing Myosin activity perturbs cell delaminations^43^, in the amnioserosa neither the number nor the timing of cell delaminations are altered (Figure 3C, E, Movie 7). These cells exhibit an accumulation of Myosin at cell junctions and a persistent, dynamic medioapical network (Figure 3I), which further accumulates in extruding cells (Figure S5). Interestingly, most of amnioserosa cells with ectopic ctMLCK show medioapical accumulation of the Rho sensor (Figure 3K), which becomes more pronounced in delaminating cells (Figure 3M, M’, M”, Movie 9). These observations, together with the junctional localization of the Rho sensor upon ectopic expression of MbsN300, show that modulating Myosin activity impacts Rho localization. ^36^

### 3) In a tissue with tension heterogeneity, cells with higher levels of MyoII are more likely to delaminate

We were intrigued by the observation that perturbing Myosin activity does not affect the overall rate of cell delaminations. While increased actomyosin contractility underlies cell ingression in several tissues^44,45^, a global increase of junctional MyoII has been found to inhibit both apoptotic basal cell delamination and apoptotic apical extrusion^43,46^. To further investigate this, we generated heterogeneities in MyoII levels across the tissue by selectively increasing or decreasing Myosin activity in groups of cells. For this, we ectopically expressed UAS-MbsN300 or UAS-ctMLCK using the prd-Gal4 driver, which is expressed in alternate segments in the epidermis and in the amnioserosa, along with the NLS:mCherry (NLS:mCh) reporter to identify MbsN300 or ctMLCK expressing cells. To assess whether changes in MyoII activity influence the likelihood of a cell delaminating, we calculated for each embryo the ratio between the number of delaminating cells NLS-mCh positive and the number of delaminating cells NLS-mCh negative.

In ECad:GFP, prdGAL4/UAS-NLS:mCh, UAS-MbsN300 embryos (Movie 11), we observe that MbsN300 expressing cells show a larger apical cell area (Figure 4A, B and Figure 6S), confirming that these cells have lower actomyosin contractility. In these embryos, the ratio of NLS-mCh delaminating cells (and thus expressing MbsN300) to non NLS-mCh delaminating cells (with wild type MyoII levels) is lower than in control embryos (Figure 4D). Conversely, in prdGAL4/UAS-NLS:mCh; UAS-ctMLCK embryos (Movie 12), we observe that ctMLCK positive cells show a smaller apical cell area (Figure 4A, C and Figure 6S). In this case, the ratio of NLS:mCh delaminating cells (with higher both junctional and medioapical MyoII) to non-NLS:mCh delaminating cells (with wild type MyoII levels) is greater than in control embryos (Figure 4D). Thus, although no increase in the number of delaminations is observed when Myosin activity is increased across the whole tissue, we find that cells with higher levels of Myo are more likely to delaminate when surrounded by cells with wild type MyoII activity. Importantly, delaminations of cells with high levels of MyoII occur throughout the entire process of dorsal closure, suggesting that high MyoII levels alone do not immediately induce delamination but increase the probability of a cell to delaminate.

**Figure 4.**
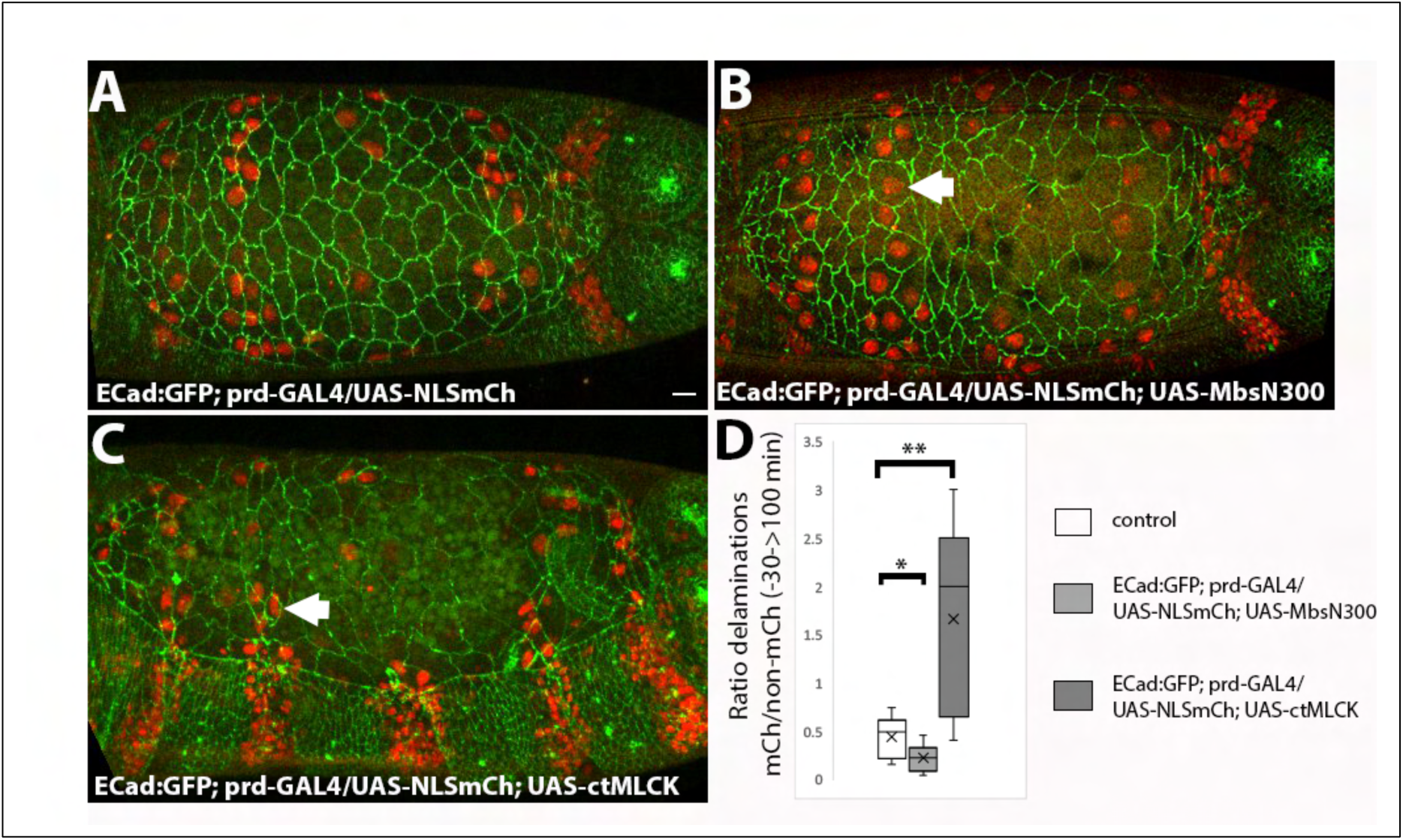
Effect of perturbing Myosin activity in groups of amnioserosa cells on cell delaminations. (A-C) Still images of from time-lapse movies of ECad:GFP; prd-GAL4/UAS-NLS:mCh (A, Movie 10), ECad:GFP; prd-GAL4/UAS-NLS:mCh; UAS-MbsN300 (B, Movie 11) and ECad:GFP; prd-GAL4/UAS-NLS:mCh; UAS-ctMLCK (C, Movie 12) embryos at 0 min of dorsal closure. (D) Ratio of NLS:mCherry to non-NLS:mCherry delaminating cells in ECad:GFP;prd-GAL4/UAS-NLS:mCh (n=11), ECad:GFP;prd-GAL4/UAS-NLS:mCh; UAS-MbsN300 (n=6) embryos and ECad:GFP;prd-GAL4/UAS-NLS:mCh; UAS-ctMLCK (n=5) embryos. Cells with lower levels of Myosin activity than their neighbours delaminate less than their neighbours (p = 0.049, Wilcoxon-Mann-Whitney test) while cells with higher levels of Myosin activity than their neighbours delaminate more (p = 0.009, Wilcoxon-Mann-Whitney test).

### 4) Blocking endocytosis prevents cell delaminations

In the pupal epidermis, a decrease in endocytic activity impairs or accelerates cell delaminations depending on the region where delaminations occur^5,29^. Therefore, we aimed to investigate the effect of blocking endocytic activity in the amnioserosa. We have previously shown that the ectopic expression of Rab5DN in the amnioserosa compromises apical constriction and apical membrane removal^47^. We observed a decrease in the number of cell delaminations with only few delaminations occurring during late stages of dorsal closure (Figure 5A, B; Movie 13).

**Figure 5.**
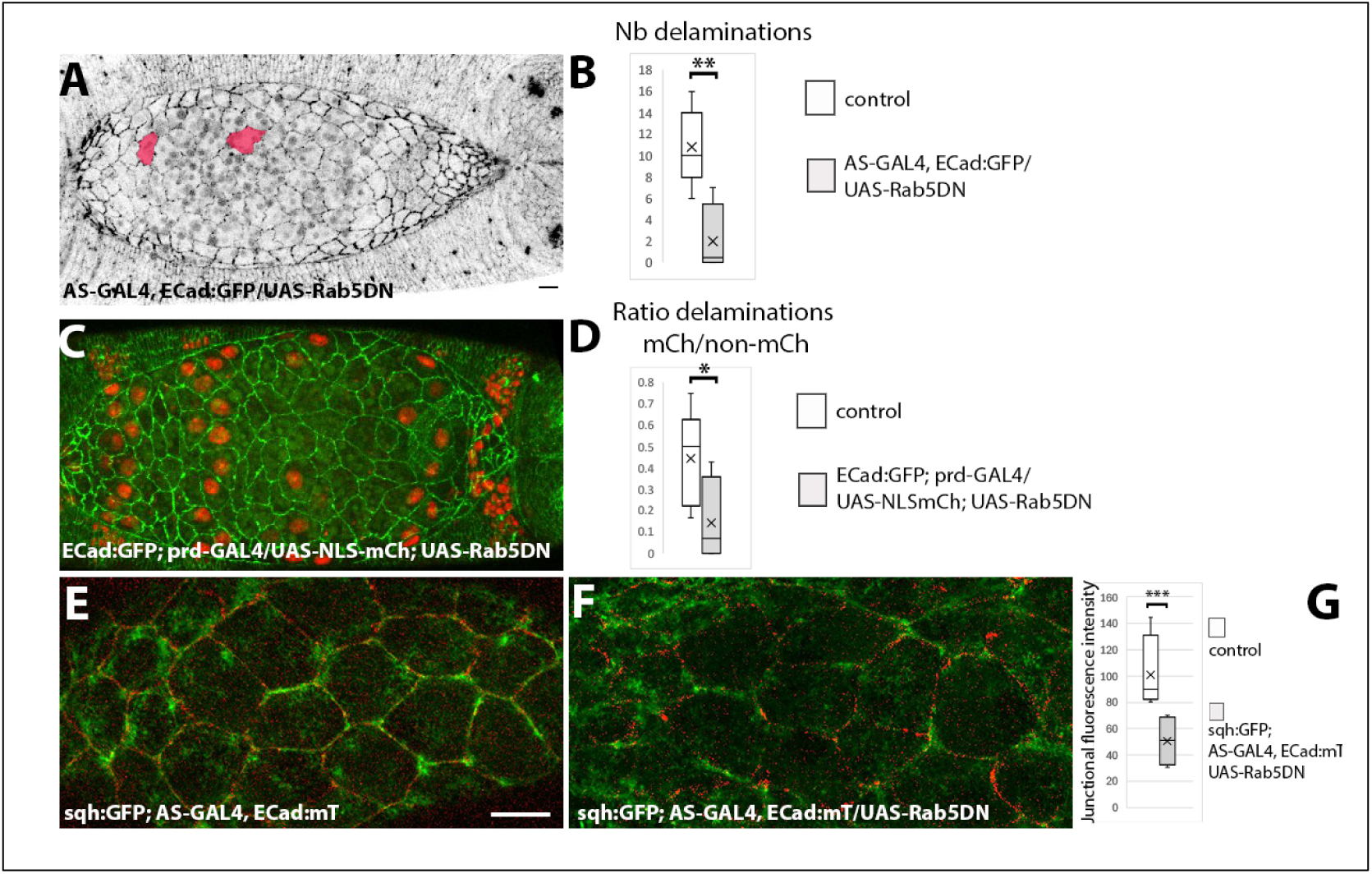
Blocking endocytosis prevents cell delaminations. (A) Still image from a time-lapse movie of an AS-GAL4, ECad:GFP/UAS-Rab5DN embryo at -30 min of dorsal closure (Movie 13). (B) Number of delaminating cells between -30 min and 100 min in control (n=7) vs AS-GAL4, ECad:GFP/UAS-Rab5DN (n=3) embryos (p < 0.001). (C) Still image from a time-lapse movie of an ECad:GFP; prd-GAL4/UAS-Rab5DN embryo at 0 min of dorsal closure (Movie 14). (D) Ratio of NLS:mCherry to non-NLS:mCherry delaminating cells in ECad:GFP;prd-GAL4/UAS-NLSmCh (n=11) and ECad:GFP;prd-GAL4/UAS-NLSmCh; UAS-Rab5DN (n=4) embryos (p = 0.03, Wilcoxon-Mann-Whitney test). (E, F) Still images from a time lapse movie from sqh:GFP; AS-GAL4, ECad:mT (E) and sqh:GFP; AS-GAL4, ECad:mT/UAS-Rab5DN (F) embryos. (G) Junctional fluorescence intensity of MyoII from still images of sqh:GFP; AS-GAL4, ECad:mT (n=4 embryos) and sqh:GFP; AS-GAL4, ECad:mT/UAS-Rab5DN embryos (n=3 embryos). Comparisons were made using a generalized mixed-effects model with embryo as a random effect and specifying a negative binomial distribution as the family (p=0.000323, generalized mixed-effects model with embryo as a random effect and a negative-binomial distribution as a family type.

The expression of Rab5DN in a mosaic manner using the prd-GAL4 driver shows that Rab5DN-expressing cells have a greater apical cell area than their neighbours (Figure 5C and S6; Movie 14), confirming that blocking endocytosis affects apical contraction. In these embryos, cells expressing Rab5DN are less likely to delaminate than control cells (Figure 5D). We examined the subcellular localization of MyoII and found that although MyoII pulses are present, there is a decrease in junctional MyoII (Figure 5E, F, G). In agreement with this observation, it has been shown that Rab5DN expressing amnioserosa cells show lower junctional tension, suggesting that endocytosis is important to maintain junctional tension^33^. These results show that endocytic activity is required for amnioserosa cell delaminations, which could be related, at least in part, to a decrease in tissue tension.

### 5) Interaction between contractility and caspase activity

Our results show that cells with increased MyoII activity due to the ectopic expression of ctMLCK are more likely to delaminate when they are surrounded by cells with control levels of MyoII. This observation raised the question of whether increased Myosin activity affects caspase activity. The apoptotic sensor Apoliner has been widely used in several systems to monitor capsase activity^48^. In this sensor, the fluorescent proteins mRFP and eGFP, are linked by a caspase-specific cleavage site from Diap1. Caspase activation leads to the cleavage of Apoliner and to the translocation of the eGFP from the cytoplasm to the nucleus. In the amnioserosa, during early stages of dorsal closure, the sensor is not active, except in the most posterior cells (Figure 6A and S7). As dorsal closure progresses, the sensor is activated first in anterior cells (Figure 6A’) and then in the whole tissue before closure is complete (Figure 6A”, Movie 15 and see also^19,37^). The activation of the sensor is completely blocked in embryos ectopically expressing Apoliner and Diap1 even after the amnioserosa has been internalized underneath the epidermis (Figure S8). Thus, Apoliner can detect low levels of caspase activity, that start to build up in the amnioserosa as dorsal closure progresses, but this activity does not inevitably lead to the cell delamination since most of the cells remain in the bulk of the tissue. We then turned to a different sensor, GC3Ai, which has been shown that can be detected in apoptotic cells throughout the whole process of programmed cell death^49^. In this sensor, the C and N termini of the GFP have been linked by a fragment containing a DEVD caspase cleavage site that must be cleaved for the GFP to fluoresce. In embryos ectopically expressing GC3Ai in the amnioserosa, sensor activity is only observed in delaminating cells (Figure 6B, Movie 16). The activity of the GC3Ai sensor in delaminating cells is detected as the cell starts to apically constrict until after it has ingressed and start to undergo fragmentation, a process that takes up to 30 minutes (Figure 6B’, B”, B’’’, B””). Thus, the GC3ai sensor detects apoptotic levels of caspase activity.

We analyzed caspase activity with both sensors in embryos with perturbed Myosin activity. We wanted to determine whether, in AS-GAL4/UAS-MbsN300 embryos, the delay in the appearance of cell delaminations was due to an absence of caspase activity or to an impairment of the expulsion of the cell from the tissue. If this was the case, we expected to see an activation of the apoptotic sensors in the tissue with no delamination. In AS-GAL4/UASApoliner, UAS-MbsN300 embryos that complete dorsal closure, the activation of the sensor followed a similar temporal pattern than in control embryos (Figure 6C, Ć, C” and S7, Movie 17), while the activity of the GC3ai sensor was only observed in delaminating cells (Figure 6E). These observations show that apoptotic levels of caspase activity do not arise independently of cell delamination.

Similarly, in embryos ectopically expressing ctMLCK, GC3Ai activity was only detected in cells that delaminate (Figure 6F). In contrast, in AS-GAL4/UASApoliner, UAS-ctMLCK embryos, we observed the presence of cells with a premature activation of Apoliner (Figure 6D, D’, D” and Figure S7, Movie 18), when comparing embryo stage using the position of the posterior spiracles. These observations suggest that increasing MyoII activity can induce low levels of caspase activity.

These observations lead us to analyze whether the preferential delamination of cells with high MyoII levels in prd-GAL4/UAS-NLS:mCh; UASctMLCK embryos was due to an increase of caspase activity induced by MyoII. For this, we asked whether delaminations of ctMLCK cells in these embryos could be prevented by the ectopic expression of an inhibitor of apoptosis. In ECadGFP; prd-GAL4/UAS-RHG miRNA; UASctMLCK embryos (where ctMLCK expressing cells were identified because of their smaller apical cell area) or in ECadGFP; prd-GAL4/UAS-NLS:mCh, UAS-RHG miRNA; UASctMLCK embryos, we didńt observe any ctMLCK expressing cell delaminating (Figure 6G, Movie 19). This result shows that the preferential delamination of cells with high MyoII cells is prevented if apoptosis is inhibited and suggests that high MyoII contractility induces cell delamination through the activation of caspase activity.

**Figure 6.**
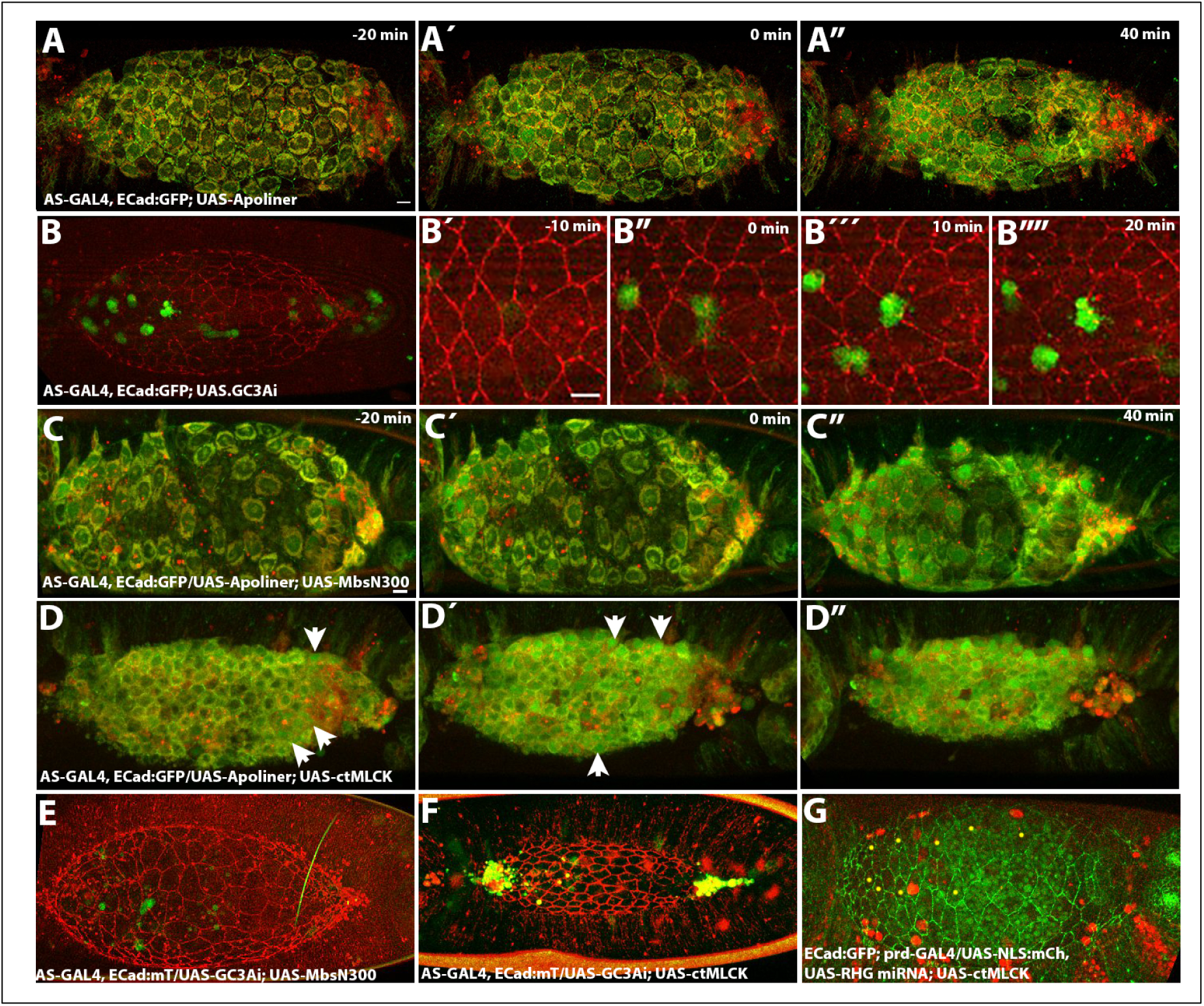
Caspase activity in the amnioserosa. (A-A”) Still images from a time-lapse movie of an example dorsal closure embryo ectopically expressing the caspase sensor Apoliner in the amnioserosa (Movie 15). Early during dorsal closure, only posterior cells show caspase activity (A,Á). As dorsal closure progresses, most amnioserosa cells located in the anterior part of the tissue start showing caspase activity while most central cells still show low levels of nuclear GFP (Á) (n=3). During late dorsal closure, all amnioserosa cells exhibit caspase activity (A”). (B) Still images from a time-lapse movie of an embryo ectopically expressing the apoptotic sensor GC3Ai (Movie 16). (B’-B””) Time course of GC3Ai activation during the delamination of an example cell based on the analysis of 20 cells from 5 embryos. (C-C”) Still images from an example AS-GAL4/UAS-Apoliner; UAS-MbsN300 embryo (Movie 17) (n=2). (D-D”) Still images from an example AS-GAL4/UAS-Apoliner; UAS-ctMLCK embryo (Movie 18) (n=6), with cells showing higher levels of caspase activity (arrows). (E, F) Still images from a time-lapse movie of an example AS-GAL4/UAS-GC3Ai; UAS-MbsN300 (F) and an AS-GAL4/UAS-GC3Ai; UAS-ctMLCK (G) embryo. Only cells that delaminate (or at the canthi) have GC3Ai signal in all the embryos analyzed (n=3 for each genotype). (G) Still image of an example ECad:GFP; prd-GAL4/UAS-NLS:mCh, UAS-RHG miRNA; UAS-ctMLCK (Movie 19). Yellow dots indicate delaminating cells. No NLS:mCh delaminating cells were observed in these embryos (n=4).

## DISCUSSION

In this work, we have investigated the mechanisms underlying the triggering and execution of apototic cell delaminations in the amnioserosa during dorsal closure, before the tissue completely ingresses into the embryo and degenerates. Our results show that cell delaminations involve, first, the stabilisation of the medioapical actomyosin cortex, and then, the formation of an actin ring, which completes the elimination of the apoptotic cell from the tissue. We have observed the appearance of actin puncta prior to the formation of the actomyosin ring, initially subapically and then co-localizing with ECadherin at the level of junctions. We propose that these puncta represent sites of actin nucleation for building the apoptotic actomyosin ring^50^. Although less prominent, Myosin puncta were also present, and we hypothesize they could be recruited by the nascent actin bundles. Our results also suggest that the small GTPase Rho regulates the reorganization of the actomyosin cytoskeleton during delamination. Rho activity has mostly been shown to be required in neighbouring cells to the delaminating cell to form a supracellular actomyosin cable that squeezes extruding cells apically from the epithelium, but its requirement in the apoptotic cell itself is more controversial^23,35^. In the amnioserosa, all delaminating cells showed accumulation of a sensor of Rho activity in the medioapical cortex, strongly supporting the role of this small GTPase in regulating the cytoskeletal changes associated with cell delamination in the delaminating cell.

Our observations do not provide evidence for the involvement of cell neighbours in the elimination of the delaminating cell. Although it has been shown that a delaminating cell induces a deformation in its neighbours and impacts their Myosin pulses^16,30^, we have not observed the formation of a supracellular actomyosin cable in neighbouring cells. An accumulation of MyoII close to cell junctions between the delaminating cell and its neighbours is only observed when ECad levels decrease significantly, suggesting that this might be a mechanism to preserve junction integrity. In fact, a mechanosensitive mechanism involving the actomyosin cytoskeleton to maintain junction integrity has been proposed^33^.

Interestingly, we find that Rho subcellular localization changes upon perturbation of Myosin activity. In embryos ectopically expressing MbsN300, both MyoII and the Rho sensor accumulate at cell junctions, while in embryos ectopically expressing ctMLCK, Myo forms a dense apicomedial mesh, and the Rho sensor shows a more persistent apicomedial localization. These observations suggests that MyoII recruits Rho either to the junctional or to the medioapical region of cells, providing evidence for the existence of a positive feedback between Rho and MyoII. We have also observed oscillatory Rho activity in non-delaminating cells (unpublished observations) and as dorsal closure progresses, medioapical Rho activity is stabilised medioapically in the whole tissue. An advection-positive feedback mechanism between Rho and its downstream effectors Rock and MyoII has been shown to underlie actomyosin pulsatility in epidermal cells during germband elongation^36^. It is possible that such a mechanism is in place in the amnioserosa, contributing to the progressive stabilization of Rho and the medioapical acomyosin cortex at the level of the whole tissue.

What triggers cell delaminations? The apoptotic removal of epithelial cells from a tissue can be regulated by both biochemical and mechanical signals^1,51^. Mechanical compression, local topological defects and the geometry of cell packing have all been shown to promote cell elimination from a tissue^31,52,53^. In the amnioserosa, it has been suggested that mechanical signals are involved in controlling cell delaminations^15,18,38^. Here, we have observed that decreasing MyoII activity in the entire tissue delays the appearance of cell delaminations. Since low MyoII activity is associated with low tissue tension^40^, our results argue for a role for mechanics in triggering cell delaminations. However, increasing tissue tension through increasing Myo levels in the whole tissue does not increase cell delaminations. A global increase in junctional Myosin has also been shown to block cell delamination in the fly notum^43^, and to disrupt apoptotic extrusion in epithelial monolayers. In the latter, increasing Myosin in apoptotic cells or in the surrounding cells disrupted extrusion^46^. This contrasts with what we observe in the amnioserosa, where cells with higher MyoII than their neighbours are more likely to delaminate. This result, together with the observation that high tissue-level Myosin activity does not affect the number of cell delaminations, suggest to us that high tissue tension or a stiff mechanical environment may exert an inhibitory effect on the ability to delaminate.

It is important to notice that cells with increased MyoII do not delaminate instantaneously but are more likely to do so over the entire process of dorsal closure. This suggests that other signals contribute to the decision of whether a cell will delaminate. In the fly notum, it has been found that the apical size of a cell, compared to other cells within the tissue and relative to the immediate neighbours, is a predictor of apoptotic cell delamination^54^. Interestingly, we have found here that cells with lower levels of Myosin and showing a greater apical cell area are less likely to delaminate, while cells with higher Myo levels and with a smaller apical cell area, are more likely to delaminate. Our results suggest that MyoII activity is a key determinant of apical cell size that could be mediating the relation between cell size and apoptotic fate. Morover, we have found that the increase in the probability to delaminate in cells with high MyoII levels is prevented upon expression of an apoptosis inhibitor, showing that high MyoII levels promote cell delaminations through the activation of caspase activity. In agreement with this, we have found that increasing Myosin activity in the whole tissue induces premature caspase activity.

The amnioserosa is a tissue in which the delamination of cells from the tissue occurs alongside the subsequent complete elimination of the tissue. This phenomenon also takes place in the larval epidermis, which is replaced by histoblasts during metamorphosis^55^, and is likely to occur in other insect extra-embryonic tissues^4^. Both in the amnioserosa and in the larval epidermis, all the cells show low, non-apoptotic, levels of caspase activity well before the tissue is eliminated. There is growing evidence that caspases have non-lethal functions and can influence a diverse array of cellular processes such as signalling, proliferation, differentiation and migration^56^. In the amnioserosa, a role for caspases in controlling cell volume decrease has been found^19^, and it is possible that low levels of caspases play a role in the remodelling of the cytoskeleton not directly linked to cell death^38^.

An important question, however, is how global tissue levels of caspases arise. We have found here that increasing Myo activity favours caspase activation. In the larval epidermis, it has also been proposed that MyoII potentiates caspase activity^5^. An interesting possibility is that actomyosin contractility influences caspase activity and thus the fate of amnioserosa cells. A relationship between mechanical cues and cell fate has been known for several years^57^. The Hippo pathway is involved in transforming mechanical stimuli into biochemical signalling pathways both in vertebrates and in Drosophila^58–60^. Notably, in wing imaginal disc cells, it has been found that medial actomyosin flows promote the medial accumulation of Kibra and the formation of a Hippo complex, which activates the Hippo pathway to repress the pro-growth transcriptional effector Yorkie^61^. The role of the Hippo pathway in the amnioserosa has not been explored yet but would be an interesting avenue for future research.

## MATERIALS AND METHODS

### Fly stocks and genetics

The following stocks were used in this work:

ECad:GFP and ECad:mT knock-in alleles^62^; sqh:GFP ^2^; aniRBD:GFP ^36^; AS-GAL4 (332.3 GAL4 line, RRID:BDSC_5398); prd-GAL4 (RRID:BDSC_1947); UAS-MbsN300^39^; UAS-ctMLCK^63^; UAS-Apoliner^48^; UAS-GC3Ai^49^; UAS-NLSmCherry (RRID:BDSC_38425); UAS-Rab5DN (RRID:BDSC_42704), UAS-Diap1 (RRID:BDSC_63820) and UAS-RHG miRNA^32^.

Stocks built during this work:

AS-GAL4, ECad:GFP and AS-GAL4, ECad:mT; sqh:GFP

AS-GAL4, ECad:mT; aniRBD:GFP

sqh:GFP; AS-GAL4, ECad:mT

UAS-NLS:mCherry; UAS-MbsN300

UAS-NLS:mCherry; UAS-ctMLCK

UAS-Apoliner; UAS-MbsN300

UAS-Apoliner; UAS-ctMLCK

UAS-GC3Ai; UAS-MbsN300

UAS-GC3Ai; UAS-ctMLCK

ECad:GFP; prdGAL4

UAS-Diap1; UAS-MbsN300

UAS-Diap1; UAS-ctMLCK

UAS-Diap1, UAS-NLS:mCherry; UAS-ctMLCK

#### Live-Imaging

Stage 12-13 Drosophila embryos were dechorionated, mounted in coverslips with the dorsal side glued to the glass and covered with Voltalef oil 10S (Attachem). The AS was imaged at 25-28°C. using an inverted LSM 710 Meta laser scanning microscope or a Leica SP8 with a 40X or a 63X oil immersion Plan-Fluor (NA=1.3) objective. GFP and mTomato cytoskeletal reporters were simultaneously imaged with an argon laser and a 651 diode laser. For whole AS imaging, 15-30 z sections 1-1.5µm apart were collected every 2 minutes. For cytoskeletal dynamics imaging, 10 z sections 1µm apart were collected every 30 seconds.

#### Image analysis and statistics

Images were processed and assembled in Fiji (version 2.16.0) and Adobe Photoshop (version 23.5.0). Images were projected via the maximum intensity projection to cover the entire tissue or a single cell in case it was in a curved region of the tissue.

Embryos were staged according to the position of the posterior spiracles: the time point at which the two contralateral posterior spiracles first meet at the dorsal side was considered the onset of dorsal closure (0 min). There was a good correspondence between the position of the posterior spiracles and the fusion of the leading edges at the anterior front of the embryo across embryos and genotypes.

The identification and tracking of delaminating cells were done manually on the maximum intensity projections and measurements of apical cell area and aspect ratio (major_axis/minor_axis of the ellipse fit) were obtained from Fiji. To identify the onset of delamination, we fitted a fifth-degree polynomial to each cell’s time-area data and computed its derivative. By scanning the derivative in reverse, we defined the onset of delamination as the point where the value first exceeded the mean.

For apical cell area measurements from amnioserosa tissues in which Myosin is perturbed in groups of cells, the tissue was segmented at 0 min of dorsal closure using Tissue Analyzer^64^ and cells with normal or perturbed levels of MyoII were identified manually.

Junctional MyoII intensity measurements were performed on maximum intensity projection of the 2-3 most apical planes identified by the ECad channel. A binary mask was created using the ECad channel which was dilated using a radius of 2 px. The mask was selected and transferred to the MyoII channel, and the fluorescence intensity of the selection was computed.

Statistical analysis of the significance of differences in phenotype across genotypes was performed using the Wilcoxon-Mann-Whitney test, linear mixed models and generalized mixed-effects models, using the R package.

## ACKNOWLEDGEMENTS

NG is very grateful to Juan José Muñoz† for his help and support during the initial phase of this project. We thank Isabel Guerrero, Julia Duque, Guy Blanchard and Luis Alberto Baena-López for critically reading the manuscript as well as Sabine Fischer for her help with fluorescence quantification and Carlos Torroja for his help with statistics. We thank the Bloomington Stock Center, Bruno Monier, Jorg Großhans and Natalia Azpiazu for Drosophila strains. Microscopy was performed at the Unidad de Citometría de Flujo y Microscopía de Fluorescencia of the Universidad Complutense de Madrid and at the Advanced Light Microscopy Facility at the Centro de Biología Molecular Severo Ochoa. This work was supported by grant PID2020-114533GB-C22 to NG from the Spanish Ministry of Economy and Competitiveness.

## Competing interests statement

The authors declare no competing interests.

## Author contributions statement

NG conceived the project, performed experiments, analysed data and wrote the paper, YF, JG and GS performed experiments, JR analysed data.

## Notes

### Competing Interest Statement

The authors have declared no competing interest.

### Summary of Updates

After receiving comments from reviewers, several changes were made in the manuscript. Specifically, some panels in Figures 1 and 6 were changed to improve clarity. Quantifications were added as well as the number of cells and embryos analyzed in each experiment. Importantly, experiment presented in Figure 6H was done using a different tool to prevent delaminations. We have now ectopically expressed RHG miRNA (three microRNAs that simultaneously inhibit the proapoptotic genes reaper, hid and grim) in the amnioserosa to prevent cell delaminations. Interestingly, we now found that delaminations are prevented in ECad:GFP; prd-GAL4/UAS-NLS:mCherry, UAS-RHG miRNA; UAS-ctMLCK embryos, in contrast to our previous result using Diap1. We believe this discrepancy arises because the effect of Diap1 is particularly sensitive to its levels of expression, something that has happened in the past in other tissues. Thus, it is possible that in our previous experiment (ECad:GFP; prd-GAL4/UAS-NLS:mCherry, UAS-Diap1,; UAS-ctMLCK embryos, which carry three UAS constructs), Diap1 levels were insufficient to fully block caspase activation and prevent cell delaminations induced by high Myosin phosphorylation. In contrast, the new experiment using UAS-RHG miRNA yields a clear result and effectively prevents the delamination of ctMLCK expressing cells. We can now conclude that the preferential delamination induced by ctMLCK requires caspase activity, and that high MyoII contractility promotes delamination through caspase activation.

