## Supplementary Information for "A role for Myosin in triggering and executing amnioserosa cell delaminations during dorsal closure"

**Figure S1**

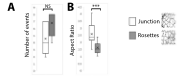

**Figure S1.** Number of delamination events for each delamination mode (A) and Aspect Ratio of cells (major_axis/minor_axis of the ellipse fit) at the onset of the delamination process for rosette vs new junction delaminations (B). No statistically significant differences were observed between the number of both delamination modes (Wilcoxon signed-rank test: W=3, p=0.3125). On the other hand, cells that leave a new junction after delamination show a significant more elongated shape than cells that delaminate forming a rosette (linear mixed model with embryo as a random effect, p=2.21e-11). Fifty-seven cells from 5 different embryos were analyzed.

**Figure S2**

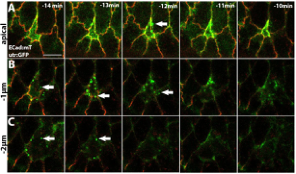

**Figure S2.** Actin dynamics during the delamination process. (A-C) Still images from a timelapse movie of a dorsal closure ECad:mT/utr:GFP embryo at the indicated time points before the completion of delamination. Apical (A, coincident with ECad:mT brightest signal) and z planes -1µm (B) and -2µm (C) basal to the apical plane are shown. Arrows indicate actin puncta. Notice the appearance of actin puncta at -2µm and -1µm, and how their localization shifts apically as apical cell area contracts.

**Figure S3**

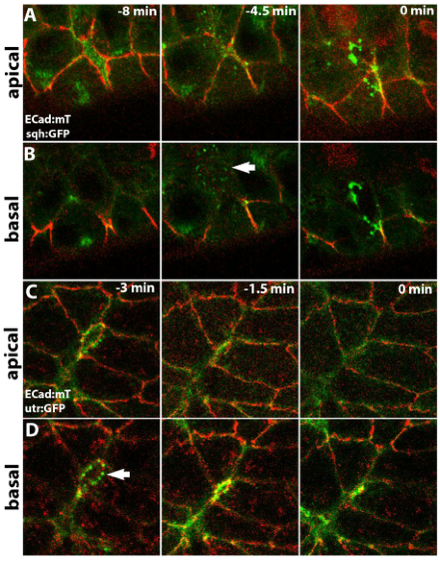

**Figure S3.** Actomyosin dynamics in cells that delaminate leaving a new junction. (A-D) Still images from a time lapse movie of a dorsal closure sqh:GFP; ECad:mT embryo (A, B) or ECad:mT/utr:GFP (C, D) at the indicated time points. Notice that the panels shown in A are maximum intensity projections of the 3 most apical planes due to the curvature of the tissue where delamination occurs; this is why in these panels (particularly at time points -4,5min and 0 min) a signal corresponding to a more basal localization is also observed. Myosin and actin puncta are indicated with arrows.

**Figure S4**

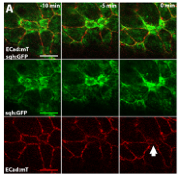

**Figure S4. (A)** Single channels of panels shown in Figure 1I, to enable visualization of ECad decrease at the level of the junctions (arrow).

**Figure S5**

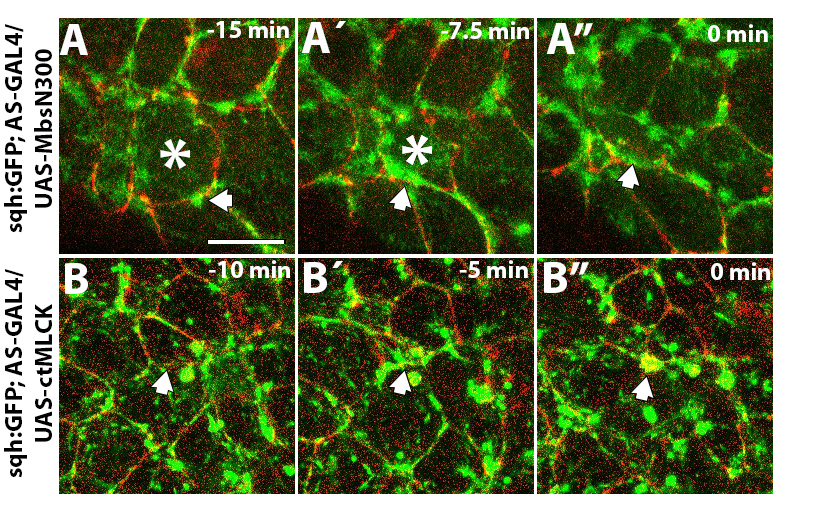

**Figure S5.** (A, B) Subcellular localization of Sqh:GFP in delaminating cells (asterisks in A and A´, arrow in B) from embryos ectopically expressing MbsN300 (A-A”) and ctMLCK (B-B”). Notice the junctional (A, arrows) and the medial (B´, arrow) accumulation of Sqh:GFP in delaminating cells. These are example cells from 5 delaminating cells from 2 embryos for each genotype.

**Figure S6**

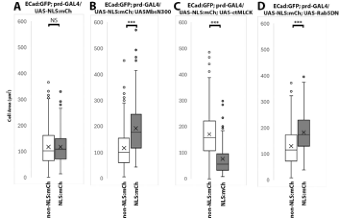

**Figure S6.** Apical cell area of NLS:mCherry and non-NLS:mCherry expressing cells at time 0 of dorsal closure for the different genotypes indicated. (A) In control embryos, no significant differences were observed in apical cell area between expressing and non-expressing cells (n= 3 embryos, number of NLS:mCherry expressing cells=151, number of non-NLS:mCherry expressing cells=359, p=0.91). (B) In ECadGFP; prd-GAL4/UAS-NLS:mCh; UAS-MbsN300 embryos, NLS:mCh expressing cells have a greater apical cell area (n=3 embryos, number of NLS:mCherry expressing cells=129, number of non-NLS:mCherry expressing cells=331, p<2e-16). (C) In contrast, in ECadGFP; prd-GAL4/UAS-NLS:mCh; UAS-ctMLCK embryos, NLS:mCh expressing cells have a smaller apical cell area (n=3 embryos, number of NLS:mCherry expressing cells=77, number of non-NLS:mCherry expressing cells=235, p=4.25e-16). (D) Finally, in ECadGFP; prd-GAL4/UAS-NLS:mCh; UAS-Rab5DNembryos, NLS:mCh expressing cells have a greater apical cell area (n=2 embryos, number of NLS:mCherry expressing cells=78, number of non-NLS:mCherry expressing cells=182, p=8.13e-07). Comparisons between NLS:mCh and non-NLS:mCh cells were done using a linear mixed model with embryo as a random effect.

**Figure S7**

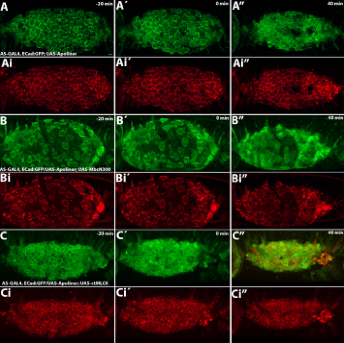

**Figure S7.** Single channels of panels shown in Figure 6A, C, D to visualize nuclear Apoliner signal.

**Figure S8**

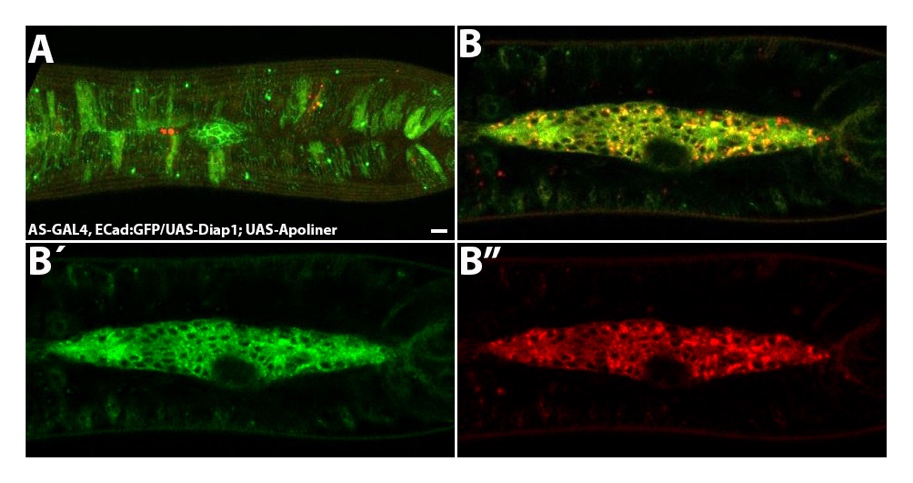

**Figure S8.** The ectopic expression of Diap1 together with the Apoliner sensor prevents activation of the sensor even after the amnioserosa has been internalized. (A) Projection of three apical planes of a late embryo showing almost complete closure. (B) Single basal plane corresponding to the same time point shown in A to show the amnioserosa that has been already internalized. (B´, B”) Single channel images to show complete absence of nuclear Apoliner localization.

**Movies:**

(Scale-bars represent 10 µm in all the movies. Unless stated otherwise, in all movies each time point is a maximum intensity projection of 20-30 z-sections 1.5µm apart and time interval is 2 minutes)

Movie 1: Time-lapse movie of an AS-GAL4, ECad:GFP embryo.

Movie 1B: This is a cropped region of previous movie isolating one delaminating cell that instead of forming a rosette, leaves a new junction.

Movie 2: Time-lapse movie of an AS-GAL4, ECad:GFP/UAS-miRNA RHG embryo.

Movie 3: Time-lapse movie of a sqh:GFP; ECad:mT embryo. Time interval is 30 seconds. Each time point is a maximum intensity projection of the 3 most apical z-zections 1 µm apart.

Movie 4: Time-lapse movie of an ECad:mT, utr:GFP embryo. Time interval is 30 seconds. Each time point is a maximum intensity projection of the 3 most apical z-zections 1 µm apart.

Movie 5: Time-lapse movie of an AS-GAL4, ECad:mT; aniRBD:GFP. Time interval is 30 seconds. Each time point is a maximum intensity projection of the 3 most apical z-zections 1 µm apart.

Movie 6: Time-lapse movie of an AS-GAL4, ECad:GFP/UAS-MbsN300 embryo.

Movie 7: Time-lapse movie of an AS-GAL4, ECad:GFP/UAS-ctMLCK embryo.

Movie 8: Time-lapse movie of an AS-GAL4, ECad:mT; aniRBD:GFP/UAS-MbsN300 embryo. Time interval is 30 seconds. Each time point is a maximum intensity projection of the 3 most apical z-zections 1 µm apart.

Movie 8: Time-lapse movie of an AS-GAL4, ECad:mT; aniRBD:GFP/UAS-ctMLCK embryo. Time interval is 30 seconds. Each time point is a maximum intensity projection of the 3 most apical z-zections 1 µm apart.

Movie 10: Time-lapse movie of an ECad.GFP; prd-GAL4/UAS-NLS:mCh embryo. Time interval is 30 seconds.

Movie 11: Time-lapse movie of an ECad.GFP; prd-GAL4/UAS-NLS:mCh; UAS-MbsN300 embryo.

Movie 12: Time-lapse movie of an ECad.GFP; prd-GAL4/UAS-NLS:mCh; UAS-ctMLCK embryo.

Movie 13: Time-lapse movie of an AS-GAL4, ECad:GFP/UAS-Rab5DN embryo.

Movie 14: Time-lapse movie of an Ecad:GFP; prd-GAL4/UAS-NLS:mCh; UAS-Rab5DN embryo.

Movie 15: Time-lapse movie of an AS-GAL4, ECad:GFP/UAS-Apoliner embryo. Time interval is 4 minutes.

Movie 16: Time-lapse movie of an AS-GAL4, ECad:mT/UAS-GC3Ai embryo.

Movie 17: Time-lapse movie of an AS-GAL4, ECad:GFP/UAS-Apoliner ; UAS-MbsN300 embryo.

Movie 18: Time-lapse movie of an AS-GAL4, ECad:GFP/UAS-Apoliner; UAS-ctMLCK embryo.

Movie 19. Time-lapse movie of an ECad:GFP; prd-GAL4/UAS-NLS:mCh, UAS-RHG miRNA; UAS-ctMLCK embryo.
